# Light stimulation system for measuring pupillary light responses based on Maxwellian view optical system using a retina presentation type viewfinder

**DOI:** 10.1101/2025.07.08.663604

**Authors:** Taisuke Eto, Takuya Yoshiike, Tomohiro Utsumi, Aoi Kawamura, Kentaro Nagao, Kenichi Kuriyama, Shingo Kitamura

**Author notes:** Corresponding authors: Dr. Taisuke Eto & Dr. Shingo Kitamura Dr. Taisuke Eto Max Planck Research Group Translational Sensory & Circadian Neuroscience, Max Planck Institute for Biological Cybernetics Adress: Max-Planck-Ring 10, 72076 Tübingen, Germany; Dr. Shingo Kitamura Department of Sleep-Wake Disorders, National Institute of Mental Health, National Center of Neurology and Psychiatry, Address: 187-1553, 4-1-1 Ogawa-Higashi, Kodaira, Tokyo, Japan.

## Abstract

**Background:** Melanopsin-expressing intrinsically photosensitive retinal ganglion cells (ipRGC) contribute predominantly to non-image-forming effects of light, such as light-induced circadian phase shift, melatonin suppression, and pupillary light reflex. Post-illumination pupil response (PIPR) has been well-used for the evaluation of ipRGC sensitivity for humans since it is able to detect ipRGC function independently from conventional cones. However, pharmacological mydriasis or a specialized light stimulation system, such as the Maxwellian view (MV) optical system, is required to avoid dynamic pupil constriction in the stimulated eye for effective PIPR measurements, which makes these approaches less practical and difficult to implement in standard experimental settings. In this context, we propose the application of a retinal projection viewfinder based on the MV system as a practical approach for measuring PIPR, and evaluate the feasibility of this approach by comparing its performance with a typical LED-based optical system.

**Methods:** Twenty-two healthy participants underwent pupillometry using both the MV-based viewfinder and a typical LED-based system. Monochromatic red and blue light stimuli were presented for durations of 1 and 10 seconds. Pupil responses, including maximum constriction, PIPR amplitude after 6 seconds from the light offset, area under the curve (AUC) values of PIPR, and sustained slopes, were analyzed using a linear mixed-effects model to assess the differences between the two systems.

**Results:** The MV-based viewfinder significantly enhanced net PIPR amplitude (*p* < 0.05) and sustained slope (*p* < 0.01) during 10-second light stimulation compared to the LED system, demonstrating its capability to effectively measure ipRGC-driven responses. In contrast, no significant differences were observed in the net AUC values. These results highlight that the MV-based viewfinder enables effective PIPR measurements by delivering constant and controlled light stimulation directly to the retina, minimizing the effects of dynamic pupil constriction during light stimulation.

**Conclusions:** The MV-based viewfinder showed feasibility as an effective method for measuring PIPR without requiring pharmacological dilation or complex optical instrumentation. This approach has strong potential for advancing the assessment of human ipRGC function using pupillary responses.

## Background

The pupillary light reflex (PLR) is an easily measurable and non-invasive physiological response. It has been widely used in various fields of research and clinical studies to investigate neural and autonomic function[1]. In particular, the post-illumination pupil response (PIPR), which represents a sustained constriction following light offset, has gained increasing attention as a non-invasive biomarker for evaluating intrinsically photosensitive retinal ganglion cells (ipRGC). These cells, discovered in the early 2000s[2–4], express the photopigment melanopsin and are the main contributors to the non-image-forming effects of light, such as light-induced circadian phase shift, melatonin suppression, and mood regulation.

Importantly, PIPR provides an approach to assess ipRGC function *in vivo* in humans in a non-invasive and relatively accessible manner, enabling the investigation of non-visual light sensitivity and its physiological consequences. In addition to the way for evaluation of basic human physiology based on ipRGC function, PIPR is being applied in studies ranging from delayed sleep-wake phase disorders[5] and seasonal affective disorders[6] to neurodegenerative diseases such as glaucoma[7,8] and Parkinson’s diseases[9]. Due to its utility in providing ipRGC-mediated responses, PIPR is increasingly used in research settings as a tool for investigating circadian biology and certain neuropathologies.

However, a significant technical challenge persists in measuring PIPR effectively. During light stimulation, the constriction of the pupil dynamically alters the amount of incident light, thereby introducing variability in the stimulus strength and impacting the reliability of measurements. A common approach to mitigate this effect is pharmacological mydriasis using a parasympathetic antagonist[10], such as tropicamide, which exerts its dilatory effects by acting on the pupillary sphincter muscle to cause its relaxation[11], but it induces glare and prolonged near vision loss lasting up to 15 hours[12]. This requires a burden on the participant to avoid driving or engaging in hazardous activities for at least the day of the mydriatic procedure[13]. The Maxwellian view (MV) optical system has been introduced as a promising solution for light delivery independent of pupil size when measuring the PIPR[14]. In contrast to the typical optical systems, where retinal illumination decreases by dynamic pupil constriction, the MV optical system focuses incident light at the plane of the pupil, thereby ensuring consistent retinal illumination that is independent of dynamic changes in pupil diameter. However, its use often requires technical knowledge and custom-built setups[15], limiting its accessibility. Recently, the MV optical system has been increasingly applied to retinal projection displays[16–18], making the system more accessible and user-friendly. Nevertheless, its applicability to pupillometry remains underexplored. Therefore, there is still a need for a more accessible, reproducible, and practically viable system for PIPR measurement that overcomes these practical and technical limitations.

To address these limitations, we propose the application of a retinal projection viewfinder based on the MV system as a practical approach for measuring PIPR, and this study aims to evaluate the effectiveness of our approach. We hypothesize that the MV-based system without the need for pharmacological dilation yields larger melanopsin-mediated metrics than the conventional LED-based system by minimizing the influence of dynamic pupil constriction, thereby bridging the gap between advanced optical setups and practical measurement needs in the field of pupillometry.

## Methods

### Participants

Twenty-two healthy adults (mean age ± standard deviation (SD): 32.7 ± 11.0 years old, age range: 19-45 years, nine females and thirteen males) participated in this study. All participants had normal color vision, as determined by the Ishihara color vision test, and no history of ocular diseases such as cataracts or glaucoma. Both oral and paper-based explanations of the study were provided to all participants before the experiment. All participants provided written informed consent to participate in this study. This study was approved by the Ethical Committee of the National Center of Neurology and Psychiatry, Japan, and was conducted in accordance with the principles of the Declaration of Helsinki.

### Equipment for light stimulation and pupil measurement

We used a retina presentation type viewfinder (RETISSA NEOVIEWER, QD Laser, Kanagawa, Japan) as a light presentation system based on the MV optical system (Fig. 1a). The three colors (red, green, blue) of a semiconductor laser and a microelectromechanical systems (MEMS) mirror are embedded in the viewfinder, and the optical paths of each color laser are regulated using two lenses and mirror (Fig. 1b)[19]. This optical configuration enables the convergence of the laser beam around the pupil, and projection directly at the retina, irrespective of pupil constriction, i.e., MV optical system. The stimuli were presented using an HDMI-connected PC that displayed RGB-filled monochromatic images (red or blue) on the 1280*720 pixels resolution display within the viewfinder. The light intensity was controlled by modulating the RGB values of the presenting image. The peak wavelengths of red and blue laser stimuli were 642 and 450 nm, respectively. As a typical light stimulation optical system, we constructed the LED-based light stimulation system (Fig. 1c). The RGB LED (WS2812B NeoPixel RGB, Adafruit Industries, New York, United States) was used as a light source, and the peak wavelengths of red and blue LEDs’ stimuli were 629 and 466 nm, respectively. The diffuser (DFB1-50C02-240, SIGMAKOKI CO., LTD, Tokyo, Japan) was located in front of the RGB LED, and a black eyecup was fixed in close contact with the diffuser for the purpose of fixing the distance from the LED to the participant’s eye and to prevent light from entering the non-stimulating eye.

**Figure 1.**
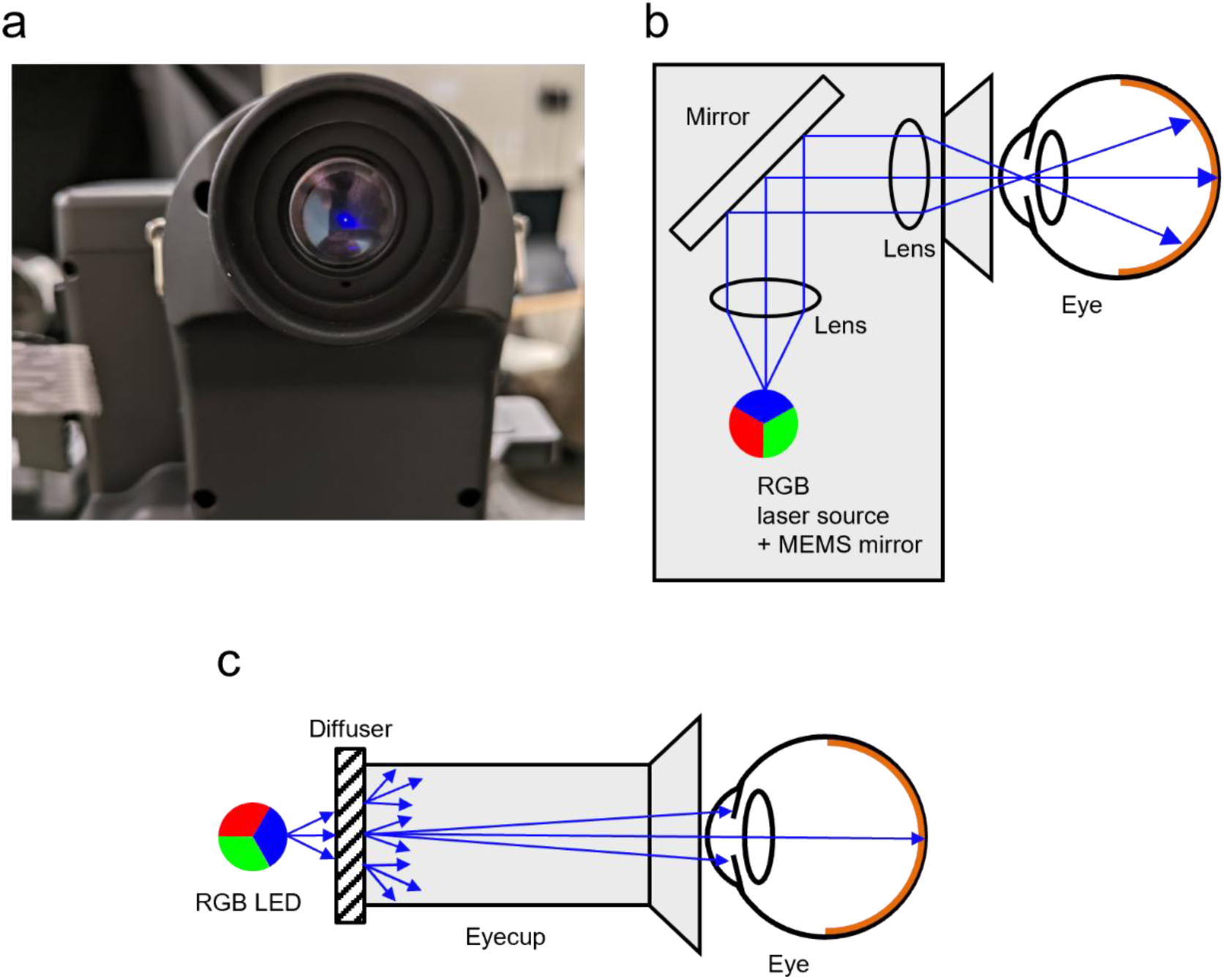
(a) An appearance of the viewfinder based on MV optical system, (c) a schematic diagram of the embedded optical system in the viewfinder and (c) a schematic diagram of the LED optical system.

The irradiance of the red and blue lasers in the MV-based viewfinder is 0.38 and 0.59 μW/cm^2^, respectively, which satisfies the Class I laser criterion set by the International Electrotechnical Commission and the International Organization for Standardization criterion (220 μw/cm^2)^. The irradiance was comparable to that of the LED-based system (0.28 and 0.43 μW/cm^2^, respectively). The photon flux and melanopic Equivalent Daylight Illuminance (EDI) standardized by the International Commission on Illumination (CIE) as CIE S 026:2018[20] of the red and blue laser were ∼12.1 log photons/cm^2^/s, 0.01 and 2.49 lx, respectively. Those were also matched to the LED-based system (∼12.0 log photons/cm^2^/s, 0.01 and 2.57 lx, respectively). All light characteristics were measured at the cornea plane of the participants using an illuminance spectroradiometer (CL-500A, KONICA MINOLTA, INC., Japan). The spectral distributions of light stimulations are shown in Fig. S1.

To measure the pupil diameter, we used a wearable glass-like pupillometer (Pupil Core, Pupil Labs GmbH, Berlin, Germany). Only the pupil of the left eye was measured while the right eye was stimulated. The sampling rate of the pupil camera was set to 60 Hz.

The MV-based viewfinder, LED system and pupillometer were controlled by a custom-built Python program based on a Python package PyPlr [21] using a Windows laptop PC. As for the LED system, an Arduino was installed between the PC and the LEDs as a relay for electronic control. When the Python program is executed with Pupil Capture, the recording software for Pupil Core, while running, a series of pupillary light response measurement protocols are performed, including measurement of the baseline pupil diameter, stimulus presentation, and post-presentation pupil measurement. For the LED system, stimuli were presented via serial communication using Arduino, and for the MV-based viewfinder, stimulus images were displayed on it by PsychoPy[22,23].

### Experimental conditions and procedures

Figure 2 illustrates the experimental protocol. All pupillometry procedures were conducted between 9:00 am and 5:00 pm to avoid circadian effects on PIPR[24].

**Figure 2.**
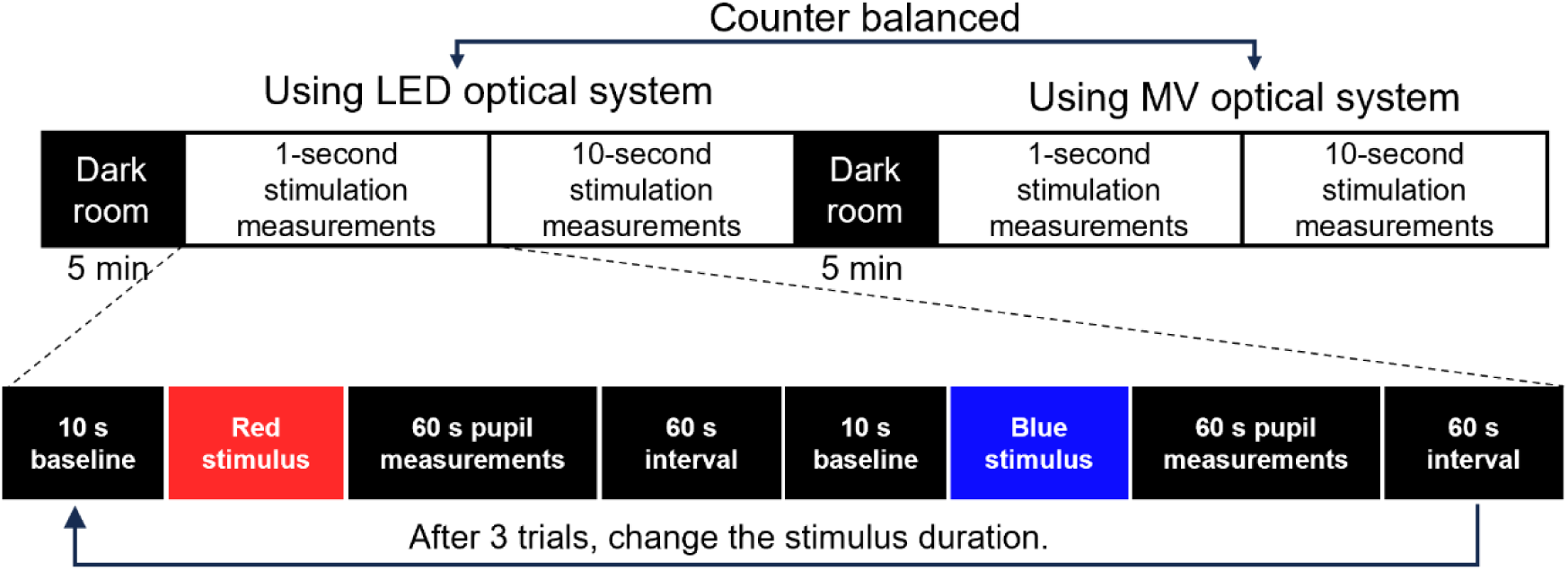
The pupil measurements experiment prodecure.

After the 5-minute dark adaptation period, participants positioned their chin on a chin rest and their right eye in an eyecup connected to the light stimulation system. Initially, pupil diameter was measured for 10 seconds in the dark environment. A long-wavelength (red) light stimulus was then presented to the right eye for 1 second, followed by a 60-second measurement of pupil diameter after the light stimulus ended. After a 60-second interval, the stimulus was switched to a short-wavelength (blue) light, and the same procedure was repeated as with the red light. This sequence (pupil measurements in the dark environment for 10 seconds, red and blue light stimuli for 1 second, and pupil measurements for 60 seconds after the light stimulus ended) was repeated three times for the 1-second light stimulation. The procedure was then repeated for the 10-second light stimulation. The above procedure was performed under both the MV-based and LED-based optical systems in a crossover manner, with the order of the two systems counterbalanced across participants. Participants were instructed to minimize blinking and to maintain fixation during stimulus presentation and pupil recording.

Pharmacological mydriasis was not used in any of the experiments, as the purpose of this study was to evaluate whether the MV system can enhance melanopsin-mediated pupil responses compared with a conventional LED system under natural pupil conditions. Best-corrected visual acuity and refractive error were not formally assessed in this study. However, previous studies have suggested that melanopsin-mediated pupil responses are independent of refractive error[25]. Furthermore, because a crossover design was used, any potential influence of refractive status would have affected both the LED and MV conditions equally and is therefore unlikely to have biased the comparison between systems.

Of the 22 participants, 10 completed only the 1-second stimulus protocol, while the remaining 12 completed both the 1- and 10-second stimulus protocol. MV-based and LED-based systems were counterbalanced. Participants were instructed to try not to blink, to keep their eyes fixed on the front, and not to move their gaze during the pupil measurements.

### Data and statistical analyses

The pupil data recorded via Pupil Capture software using a custom-built Python program was converted by the data export software Pupil Player for Pupil Core. The converted pupil data were initially analyzed using a Python program based on PyPlr[21], and the pupil diameter was converted to a relative pupil diameter [%] based on the pupil diameter in the dark environment for 3 seconds before the stimulus was presented. Blink-related artifacts were automatically detected and handled using the preprocessing functions implemented in Pyplr.

All the following analyses were performed using R version 4.4.0 (R Core Team, Austria)[26]. Relative pupil diameter data were applied noise rejection using the Savitzky-Golay filtering with “signal”[27] package as well as PyPlr. The pupil response variables: (1) Maximum pupil constriction, (2) PIPR amplitude, (3) AUC early, (4) AUC late, and (5) Sustained slope during light stimulation, were extracted from this smoothed data according to previous studies[15,28,29]. The maximum pupil constriction was defined as absolute values when the relative pupil diameter was at its minimum due to light stimulation, and the PIPR amplitude was defined as the mean absolute values of relative pupil diameter between 5.5 and 6.5 seconds after light offset. The AUC early and AUC late were calculated using the R package “pracma”[30] as a value of a trapezoidal numerical integral of the relative pupil diameter between 2 and 10 seconds after light offset and 10 and 30 seconds after light offset, respectively. The sustained slope was defined as a slope of best-fit linear regression from 2 to 10 seconds after light onset in the only 10-second stimulation condition. Net values were calculated by subtracting red from the value for variables (1) – (4) and by subtracting blue from red for variables (5) so that the increasing net value indicates more active ipRGC responses.

All statistical analyses were also conducted using R version 4.4.0 with R packages with “lmerTest”[31] and “emmeans”[32] to perform the linear mixed-effects model (LMM) and calculate the estimated marginal means (EMMs), respectively. The LMMs were applied to all “net” PLR and PIPR variables except the sustained slope, with “condition (MV or LED)” and “duration (1 or 10-second)” as fixed effects, and “subject,” “number of trials,” “age” and “sex” as random effects. These models took into account the interaction effects of “condition” and “duration.” For the sustained slope, since the stimulus duration was only 10 seconds, we conducted an LMM that excluded “duration” from the fixed effects.

## Results

Three participants were excluded: one withdrew at the participant’s request, and two exhibited excessive sleepiness during the measurements. Of the remaining 19 participants included in the final analysis, 11 completed the 10-second stimulation protocol (mean age ± standard deviation [SD]: 31.2 ± 12.0 years old, three females and eight males), whereas all 19 completed the 1-second protocol (mean age ± SD: 33.2 ± 11.3 years old, eight females and eleven males).

The mean relative pupil diameter (%) was shown for each light color and optical system condition in 1 (Fig. 3a) and 10-second stimulation (Fig. 3b). To verify that pupil diameter had returned to baseline before the next trial, we calculated the difference between the mean pupil diameter during the baseline period (−3 to 0 s) and that during the final 3 s of the recording for each trial. A LMM with subject as a random effect showed no significant deviation from zero (estimate = −1.42 %, *p* = 0.103), indicating that pupil diameter had returned to baseline before the next stimulus.

**Figure 3.**
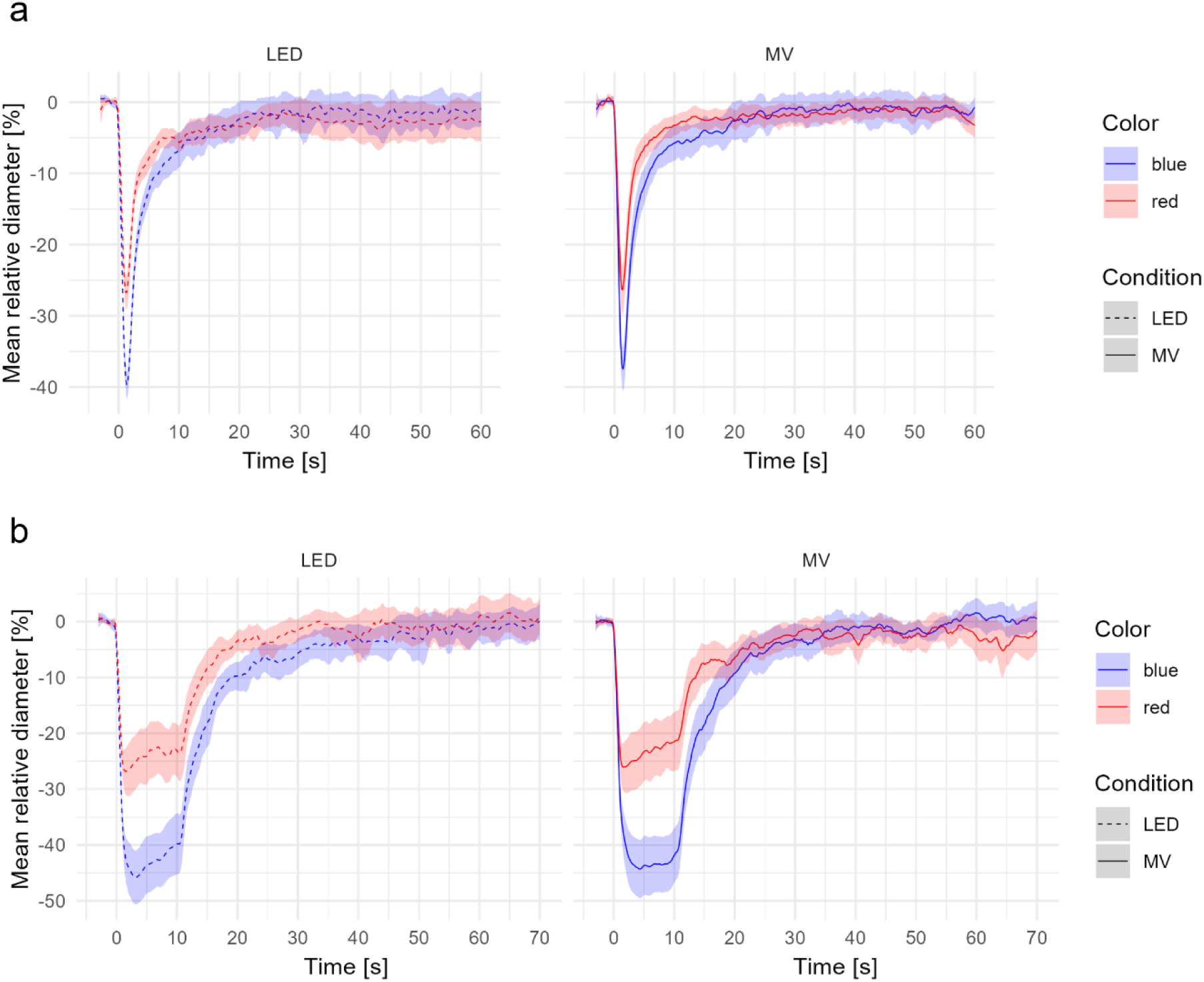
The mean relative pupil diameter [%] for (a) 1-second stimulation and (b) 10-second stimulation. The red line and red shaded area represent the mean value and standard error of the mean (SEM) for the red light stimulation, respectively. The blue line and blue shaded area represent the mean value and SEM for the blue light stimulation, respectively. The dashed and solid lines present LED and MV conditions, respectively.

Figure 4 shows the distribution of the net pupil response variables for each optical system condition and stimulation duration. Each dot represents an individual recording obtained from repeated measurements of each participant. For the (1) net maximum pupil constriction, the LMM revealed a significant main effect of stimulation duration (*t*(169.5) = 2.47, *p* = 0.015), indicating that the larger net maximum pupil constriction in 10-second stimulation (estimate = 4.09, standard error: SE = 1.66) than that in 1-second stimulation. On the other hand, no significant effects were observed for optical system conditions (*t*(157.4) = −1.2, *p* = 0.23) and interaction effects of conditions and stimulation duration (*t*(157.1) = 0.13, *p* = 0.90) (Fig. 4a).

**Figure 4.**
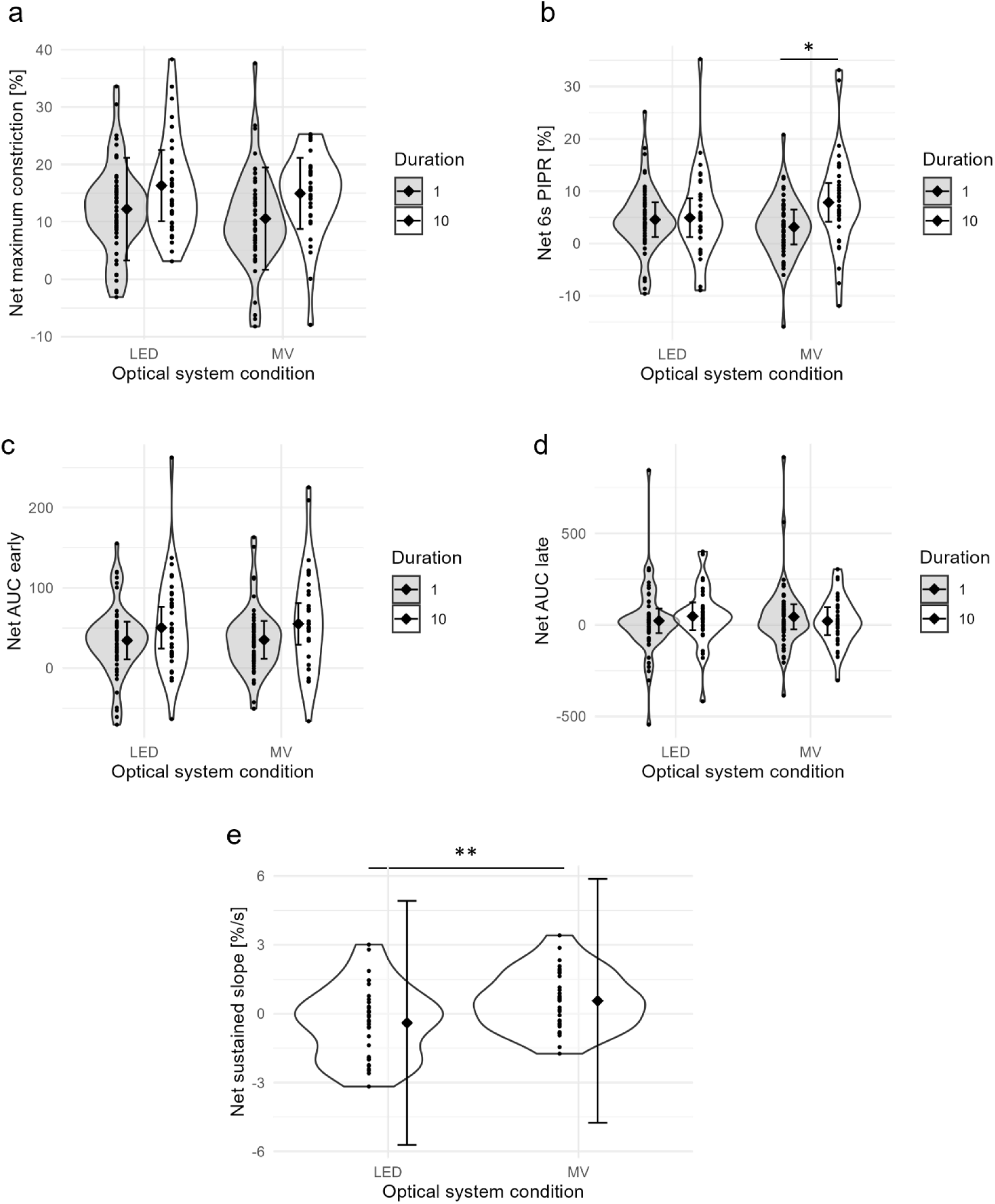
The violin plots of the net pupil reponses valuables: (a) net maximum constriction, (b) net PIPR amplitude around 6 seconds after the light offset, (c) net AUC early, (d) net AUC late and (e) net sustained slope. Each dot represents an individual recording obtained from repeated measurements of each participant. The diamond shape and error bar represent the estimated merged mean and standard error of the mean. *: *p* < 0.05, **: *p* < 0.01.

For the (2) net PIPR amplitude around 6 seconds after the light offset, no significant main effect of optical system conditions (*t*(159.7) = −1.14, *p* = 0.27) and stimulation durations (*t*(166.6) = 0.25, *p* = 0.81) were found, meanwhile a significant interaction effect of optical system conditions and stimulation durations was observed (*t*(158.4) = 2.08, *p* = 0.039). The net PIPR amplitude was significantly higher in the MV condition and 10-second stimulation (estimate = 4.32, SE = 2.08, *p* = 0.018) than that in the MV condition and 1-second stimulation (Fig. 4b).

Neither the (3) net AUC early (Fig. 4c) nor (4) net AUC late (Fig. 4d) was significantly affected by the optical system conditions (AUC early: *p* = 0.93, AUC late: *p* = 0.70), stimulation duration (AUC early: *p* = 0.11, AUC late: *p* = 0.72) and interaction of the conditions and duration (AUC early: *p* = 0.59, AUC late: *p* = 0.46).

For the (5) net sustained slope during 10-second light stimulation, a significant effect of the optical system conditions was shown, and the net sustained slope in the MV condition was significantly larger than that in the LED condition (*p* = 0.0054, Fig. 4e).

## Discussion

This study investigated the feasibility of using a retinal projection-type viewfinder based on a Maxwellian view (MV) optical system to measure ipRGC-mediated pupil responses, particularly the post-illumination pupil response (PIPR). By comparing this system with a conventional LED optical system, we examined several key pupil response variables, namely maximum constriction, PIPR amplitude, AUC early, AUC late, and sustained slope during light stimulation. We found that the MV-based system marginally but significantly enhanced PIPR amplitude and sustained slope during light stimulation. These findings partially supported that this MV system could be applied to effectively measure the ipRGC-derived pupil response rather than the conventional LED system only under a 10-second condition, whereas 1-second responses are comparable. To our knowledge, this is the first study to quantify PIPR using an integrated, head-mounted Maxwellian-view display, extending classic bench-top MV work[14].

For the results of the net PIPR amplitude around 6 seconds after the light offset under 10-second light stimulation, the larger net PIPR amplitude obtained with the MV-based system further support its utility of this system, with the weak irradiance (∼12.1 log photons/cm^2^/s) which is comparable to a laser safety criteria, for application to PIPR measurement. This corroborates the irradiance-response trend reported by Adhikari et al[33]. With a 1-second duration, the attenuation of light intensity due to pupil constriction may not contribute significantly to the PIPR. In particular, the light intensity of stimulation in this study was very weak at around 2 lx in mEDI, and even with 50% pupil constriction, the light intensity would only be attenuated by at most 1 lx with mEDI. On the other hand, in the case of 10-second light stimulation, a cumulative difference in intensity of the light reaching the retina is likely to lead to greater ipRGCs activation in the MV condition. Because the MV optics project the stimulus through a nearly fixed entrance pupil (∼1 mm), the retinal irradiance remains stable despite a 30-50 % pupillary constriction, whereas the LED system loses up to ∼ 1 log unit over 10 s. This may explain the larger cumulative melanopsin drive under the MV system. The larger net sustained slope during 10-second stimulation further supported this interpretation. This enhancement of melanopsin-mediated effects with longer stimulus duration is corroborated with the known characteristics of ipRGCs, such as sustained PIPR responses in duration-dependent manner and the role of melanopsin in prolonged pupil response with linger light exposure[33,34]. Although the spectral composition and irradiance slightly differed between the LED and MV systems, the melanopic EDI was matched between conditions. Therefore, the effective stimulus strength for melanopsin photoreception was comparable across the two systems.

In the net AUC early and late, no significant effects of the optical system conditions, stimulation duration, and interaction between the two were observed. It has been reported that the intra- and inter-individual differences for the net AUC early and late are highly variable, especially at low light intensities[33]. The measurement instability may disrupt the potential effects of each factor, i.e., stimulation duration and optical system conditions. The benefits of the MV optical system may be highlighted in future work with higher light intensity.

Although this study demonstrates the feasibility of applying the retinal projection viewfinder based on the MV system to the PIPR measurement, it has several limitations. First, the sample size was relatively small, especially in the 10-second stimulation condition (n = 11). Second, the intensity of light stimuli was weak (12.0 log photons/cm^2^/s) due to the safety limitations of the laser compared with the generally used light intensity in PIPR measurements[6–8,29]. Nevertheless, it has been shown that the ipRGC is activated >11 log photons/cm^2^/s irradiance[35], and indeed, ipRGC-derived sustained pupil responses have been observed in this study. However, the relatively low stimulus intensity may have limited the ability to detect differences between optical systems under the 1-s stimulation condition. In contrast, during the 10-s stimulation, the cumulative effect of light exposure may have allowed the advantage of the MV system to become detectable. Future studies should investigate whether higher stimulus intensities would enable similar differences between systems to be observed even with shorter stimulation durations, which would be preferable for practical applications due to the reduced burden on participants.

### Conclusion

In conclusion, we demonstrated the applicability of the retina presentation type viewfinder based on the MV optical system for assessing ipRGC-mediated pupil responses to the light, i.e., PIPR. As the development of retina presentation-type displays and viewfinders has been actively conducted[16–19], effective PIPR measurement using the MV optical systems may become a useful tool for research and potential clinical applications, and this study provides an initial demonstration of its feasibility. The findings suggest that the proposed pupil response measurement system may contribute to the development of non-invasive approaches for investigating ipRGC function and related physiological processes.

## Declarations

### Ethics approval and consent to participate

This study was approved by the Ethical Committee of the National Center of Neurology and Psychiatry, Japan. All participants provided written informed consent to participate in this study.

### Consent for publication

Not applicable.

### Availability of data and materials

The datasets used and/or analysed during the current study are available from the corresponding author on reasonable request.

### Author’s contributions

**TE**: Conceptualization, Methodology, Software, Validation, Formal analysis, Investigation, Resources, Data Curation, Writing – Original Draft, Visualization, Project administration, Funding acquisition; **TY**: Writing – Review & Editing; **TU**: Writing – Review & Editing; **AK**: Writing – Review & Editing; **KN**: Writing – Review & Editing; **KK**: Resources, Writing – Review & Editing; **SK**: Conceptualization, Methodology, Resources, Supervision, Project administration, Funding acquisition.

## Acknowledgments

The authors declare that they used ChatGPT (GPT-4o, OpenAI) to assist with English language editing and refinement of this manuscript. The final content was reviewed and approved by all authors.

## List of abbreviations

ipRGC: intrinsically photosensitive Retinal Ganglion Cell
MV: Maxiwellian view
PIPR: post-illumination pupil response
AUC: area under the curve
LMM: lenear mixed-effects model

